# Lipid Organization by the Caveolin-1 Complex

**DOI:** 10.1101/2024.07.10.602986

**Authors:** Korbinian Liebl, Gregory A. Voth

**Author notes:** **Corresponding Author:** Gregory A. Voth, Department of Chemistry, The University of Chicago, 5735 S. Ellis Ave, SCL 123, Chicago, IL 60637, **E-mail:**.

## Abstract

Caveolins are lipid-binding proteins that can organize membrane remodeling and oligomerize into the 8S-complex. The CAV1 8S-complex comprises a disk-like structure, about 15nm in diameter, with a central beta barrel. Further oligomerization of 8S-complexes remodels the membrane into caveolae vessels, with a dependence on cholesterol concentration. However, the molecular mechanisms behind membrane remodeling and cholesterol filtering are still not understood. Performing atomistic Molecular Dynamics simulations in combination with advanced sampling techniques, we describe how the CAV1-8S complex bends the membrane and accumulates cholesterol. Here, our simulations show an enhancing effect by the palmitoylations of CAV1, and we predict that the CAV1-8S complex can extract cholesterol molecules from the lipid bilayer and accommodate them in its beta barrel. Through backmapping to the all-atom level we also conclude that the Martini v2 coarse-grained forcefield overestimates membrane bending, as the atomistic simulations exhibit only very localized bending.

## Introduction

The Caveolin-1 (CAV1) protein consists of 178 amino acids and oligomerizes into the CAV1-8S complex. The structure of the CAV1-8S complex has been resolved recently, showing that eleven protomers assemble into a disk-like structure ∼15nm in diameter with a central beta barrel (1-3). Importantly, each protomer of CAV1 contains three palmitoylation sites (4-6), and the CAV1-8S complex is monotopic, i.e., it is embedded only in the outer leaflet of the membrane (3,7). Structural data suggests a flat-membrane exposed surface of the CAV1-8S complex. Clustering of multiple complexes gives rise to caveolae biogenesis, hypothetically by inducing a cholesterol dependent remodeling of the global membrane into a dodecahedral shape with the CAV1-8S complexes located on the flat polyhedral areas (2,8-12).

Its substantial capability for sensing tension and remodeling of the membrane empowers the CAV1-8S complexes for many biological roles (13,14). Amongst others, CAV1 regulates endocytosis, cell signaling and the lipoprotein metabolism. Dysfunction of CAV1 is hence related to several diseases including lipodystrophy, cardiac hypertrophy and diabetes (13,15-18). Expression of CAV1, on the other hand, plays dual roles in cancer, where it can act as both a tumor suppressor and promoter (19,20). In spite of the high biological relevance, many of CAV1’s molecular mechanisms have remained poorly understood. The role of its palmitoylations is one major open question for CAV1. Dietzen et al and Frank et al have reported that palmitoylation of CAV1 is in principle not required for caveolae formation, though it may affect affinities (4,21). Koh et al, in turn, have emphasized that palmitoylations of CAV1 are important for proper synaptic functioning. Elimination of palmitoylations (via mutation of the associated cysteines) substantially alters synaptical vesicle endo- and exocytosis, for example (22). Also the mechanism behind membrane remodeling during caveolae formation is largely unexplained. It is understood that dodecahedral caveolae are stabilized by Cavin proteins likely due to electrostatic and entropic interactions, but the initial flat to curved transition is generated through CAV1 complexes by an unexplored mechanism (23,24).

Moreover, one may raise the question how the behavior described above correlates with cholesterol binding and accumulation. Since these open questions are molecular in nature, Molecular Dynamics (MD) and coarse-grained (CG) simulations represent attractive methods to address them. Interactions between lipid bilayers and a single CAV1 protein have been studied extensively by the Sengupta group using Coarse-grained simulations. These simulations indicate that membrane bending and cholesterol accumulation occurs coincidently and suggest that the palmitoyl tails may increase binding affinity of CAV1 to the membrane, whereas the membrane composition itself is not significantly affected by it (25,26). However, these results rely on the description by the Martini model and concern CAV1 monomers. Thus, the interpretations are not simply transferable to the recently resolved CAV1-8S complex, leaving its molecular mechanisms elusive further on.

For these reasons, we have performed extensive atomistic Molecular Dynamics (MD) simulations of the CAV1-8S complex embedded in a lipid bilayer with 30% Cholesterol content and exposed to explicit solvent yielding system sizes of ∼1 million atoms. Simulations were performed comparatively, with and without palmitoylation tails, for up to 1 µs. These unbiased simulations indicate distinct cholesterol coordination in the presence of palmitoylation, but we emphasize that the simulation timescale for such a large system is still too low for effectively sampling complete reorganization of the membrane.

In order to generate broader sampling of protein-lipid compositions, we have therefore also performed well-tempered Metadynamics simulations using a coordination number as the collective variable (27,28). The substantially larger conformational space covered with this method manifests itself by indicating a free energy minimum that has not been sampled at all during the unbiased 1µs simulations. Thus, the Metadynamics simulations provide several novel molecular insights. For instance, we find that the CAV1-8S complex can extract individual cholesterol molecules from the inner membrane layer and accommodate them in its beta barrel. This mechanism occurs independent from the palmitoylation state. Otherwise, the Metadynamics simulations show that palmitoylations organize an ordered, vertically stacked cholesterol packing in the inner layer and enhance cholesterol accumulation.

Although our overall simulation methodology enables the sampling of higher cholesterol concentration, we did not find significant changes in membrane bending during the unbiased simulations. We note, however, that Martini CG simulations we will describe below are more indicative of stronger membrane bending. In order to explore this prediction, we have backmapped a CG structure from a Martini v2 simulation to an atomistic structure and performed an atomistic simulation starting from the backmapped structure. Throughout the resulting all-atom simulation, the membrane-bending decreased in contrast to the Martini prediction. Thus, our simulations suggest that the flat-to-curved transition for caveolae arises as a more global, cooperative effect generated by several CAV1-8S complexes. We note that the CG Martini forcefield has a number of physical issues, one of which is an incorrect description of the “missing entropy” in the CG model and another of which is an anomalous degree of bending (undulation) of membranes in the intermediate lengthscale regime (29).

## Method and Materials

### System preparation and unbiased simulations

The 7SC0 pdb-structure was used as model structure for the CAV1-8S complex (3). This structure was processed with the *PPM* webserver to determine positioning of the complex in the lipid bilayer (30). Consistent with experimental findings, monotopic orientation of the protein complex was found. The output was used in subsequent system preparation with *CHARMM-GUI* (31,32). Here, two distinct systems were generated: One entirely lacking palmitoylation, and one with S-palmitoylation of the 133, 143 and 156 Cysteine residues of each monomer (in total 33 S-palmitoylated cysteines). Lipid bilayers, consisting of 70% POPC and 30% Cholesterol, with 31.7 nm in x- and y-dimension were built, and the Charmm36m force field was employed (33). The systems were solvated with the TIP3P water model and a salt concentration of 150 mM NaCl was adjusted on top of electrostatic neutralization (34). The z-dimension of the systems were 13.3 nm. Energy minimization was carried out for at least 10000 steps using the steepest-descent method with the Gromacs/2020 package, which was also used for all subsequent MD simulations (35). In cases where the energy minimization crashed due to too strong atomistic overlap, we used a Gromacs version compiled at double-precision. Afterward, the systems were equilibrated according to the output generated by *CHARMM-GUI*. In the first of two stages, systems were simulated in the constant NVT ensemble using the Berendsen thermostat at a reference temperature of 310.0 K and a coupling constant of 1 ps (36). In the following four equilibration simulations, a semi-isotropic Berendsen barostat was used with a reference pressure of 1 bar, coupling constant of 5 ps and compressibility 4.5e-5 bar^-1^ (36). Conventional restraints were used and sequentially reduced in the equilibration phase that lasted 1125000 steps. The first 375000 steps were propagated by 1 fs and the remaining steps by a time-step of 2 fs, yielding a total equilibration time of 1.875 ns. The output structure was used for the production runs that were performed for 1µs using an MD time-step of 2fs. In the production runs, no restraints were turned on and the Noose-Hoover thermostat and Parrinello-Rahman barostat were employed with same coupling and reference constants as in the equilibration phase (37-39). Simulations were analyzed with the MDAnalysis package and VMD was used for visualization (40-42).

### Metadynamics simulations

The well-tempered Metadynamics simulations were performed using Gromacs in combination with the PLUMED package (27,43,44). The sum over all contacts (eq. 1) of C27 Cholesterol- atoms (end of hydrocarbon sidechain) with the CZ atoms of the PHE167 residues (in the beta-barrel) was used as reaction coordinate. The *r*_0_ parameter was set to 1.5 nm, d_0_ to 0.0 nm, and neighbor list updating with a cutoff of 10 nm was activated. In the well-tempered Metadynamics simulations, the reaction coordinate was biased by adding Gaussian potentials every 1000 steps. Gaussian potentials had a width of 2.5 (a.u.) and a height of 0.5 kJ/mol scaled with a bias factor of 15 and a temperature of 310 K. Grid spacing was applied with 150 bins. Metadynamics simulations were performed for ∼ 220 ns.

### Martini simulation

The preparation and equilibration for a Martini simulation was conducted similarly to the atomistic systems using the *Martini Maker* in *CHARMM-GUI (31*,*45*,*46)*. The box-dimensions were 35.0,35.0,12.5 nm in x,y,z direction. The lipid composition was 30% cholesterol and 70% POPC. Palmitoylation was not included, and the system was described by the Martini 2.2 model (47). Note that a cholesterol-description compatible with Martini 3 was released when we were already finalizing our work (48,49). However, the Martini 3 version is still limited by entropy-enthalpy compensation(48). Accurate description of the CAV1-8S complex may hence still be a challenge, that is likely to be addressed in a future project.

For the 100ns production run, a time-step of 5 fs (instead of 20 fs) was used. Velocity rescaling was used as thermostat ensuring a reference temperature of 310 K and a coupling constant of 1 ps (50). Semiisotropic pressure-coupling with the Parrinello-Rahman barostat was applied with a coupling constant of 12 ps, and a compressibility of 3.4e-4 bar^-1^ at a reference pressure of 1 bar. Further specific settings were nstpcouple=1, rlist=1.9, nstcalcenergy=1, nsttcouple=1 and nstlist=1.

### Backmapping

As a first attempt to obtain an atomistic structure mapped back from a CG Martini structure, the Martini to All-Atom Converter integrated in *CHARMM-GUI* was used (31,51). The resulting output structure, however, showed numerous cholesterol-ring penetrations by the POPC tails. These artifacts could not be resolved by energy-minimization. Thus, the recently developed Multiscale Simulation Tool (mstool) was instead applied to backmap the lipid bilayer, as mstool has been successfully used for these systems (52). The backmapped bilayer structure of our system captures the membrane-shape accurately, and no cholesterol-ring penetrations in the output-structure were obtained. The final backmapped structure was processed analogously to the standard atomistic simulations, whereby the same box extensions (x,y: 42.6 nm, z: 27.5 nm) of the Martini structure were adjusted for the atomistic simulation. The solvated system was energy minimized and equilibrated as outlined in the *System preparation and unbiased simulations* section. Afterwards, a production run was performed for ∼360ns.

## Results

### Unbiased atomistic simulations

During the 1 µs MD simulations, we observed that the disk-like structure and central barrel of the CAV1-8S complex remains stable with and without palmitoylation tails (Fig 1). Fluctuations described by the rmsd-curves are slightly elevated by partial unfolding of the N-termini, which nevertheless remain contactless with the lipids. Overall, we recorded only modest changes in lipid structure and composition on this timescale, and the CAV1-8S complex remained monotopic with only a minor vertical protrusion from the bilayer occurring during the first half of the simulation (measured as mean distance between phosphate atoms of upper layer and the residues 85-86 of all monomers, Fig 1D). As shown in Fig 1 E, the number of cholesterols at the central beta-barrel slightly increases throughout the simulation for both systems, with and without palmitoylation. However, diffusion of lipids is small on the timescale of a few microseconds. Longer timescales are not readily reachable for our atomistic MD systems that consist of ∼1 million atoms.

**Figure 1:**
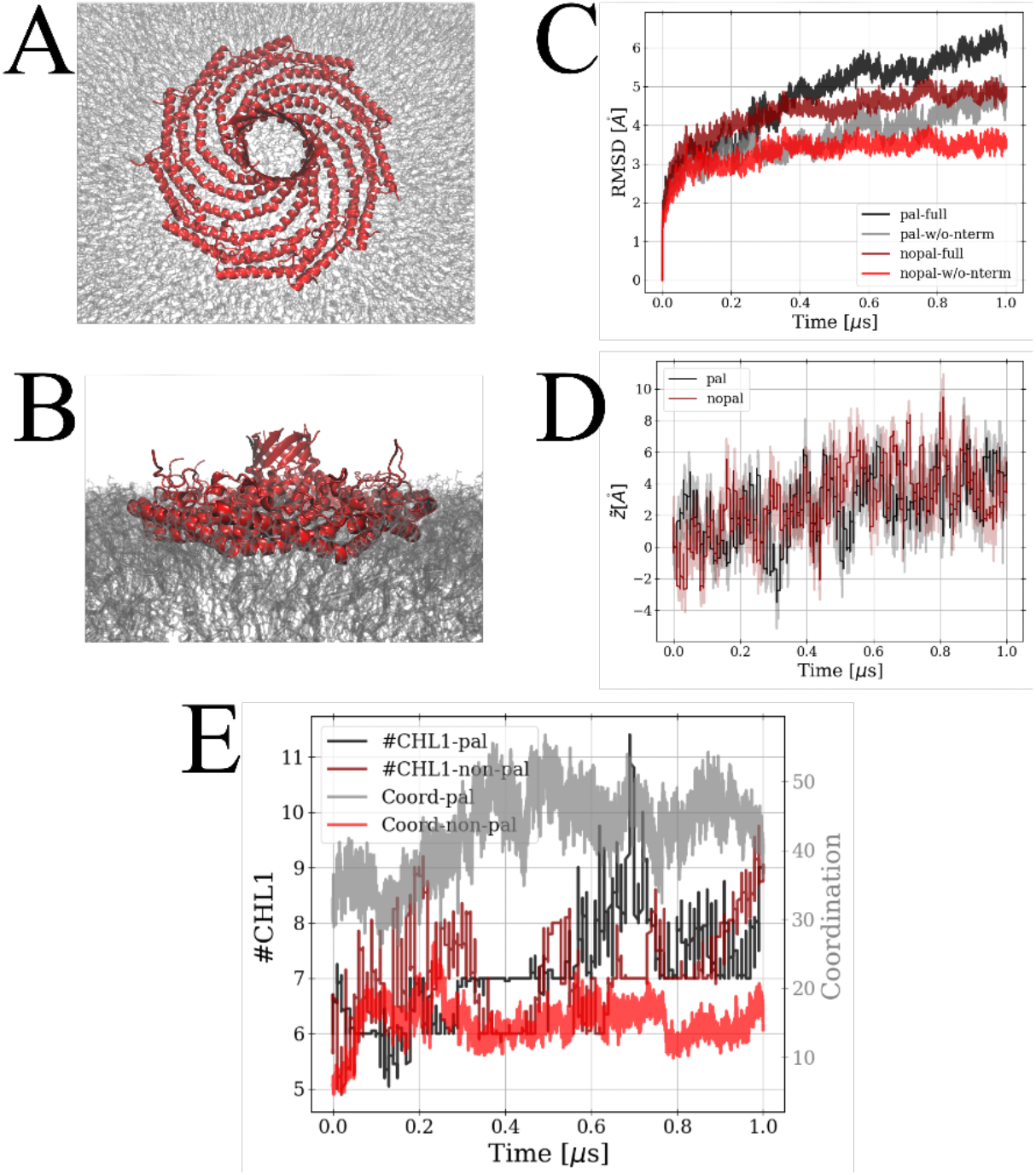
Structural snapshots of the CAV1- 8S complex (red) embedded in the lipid bilayer, from top perspective (A), and side-view (B). Root-mean square deviations during the simulations have been calculated including and excluding the N-terminal amino acids, respectively (C), and minor changes in protein height relative to the lipid bilayer are noted (D). The number of cholesterol molecules within a cylindrical volume of 1.5 nm radius centered at the protein complex are shown as thin lines (black for palmitoylated and darkred for unmodified system) in (E). The postprocessed coordination number (eq 1) is plotted as thicker lines (grey and red, respectively).

For these reasons, we designed an advanced sampling setup that employs a collective variable between cholesterol and atoms of the CAV1-8S beta-barrel. The coordination number is the sum over all contacts s_ij_ between specific cholesterol and protein atoms:

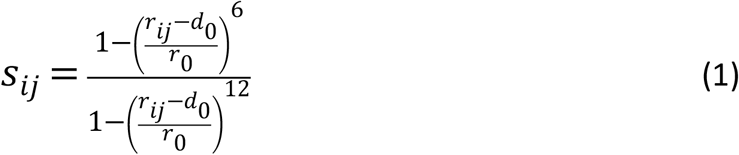

Details on this setup are given in the Methods section. Note that the coordination number directly correlates with the number of cholesterol molecules accumulated at the central beta barrel; correlation coefficients are 0.48 for the palmitoylated and 0.34 for the unmodified system, respectively. However, this also depends on the orientation of the cholesterol molecules. Thus, the higher coordination number sampled in the palmitoylated case indicates that the palmitoyl tails regulate the orientational alignment of cholesterols. We used the above coordination number for subsequent well-tempered Metadynamics simulations, and we limit here our further analysis to those simulations, as we find considerably better exploration of the phase space in these simulations.

### Metadynamics simulations

As revealed by the rapid influx of cholesterol molecules toward the central beta barrel of the CAV1-8S complex, the well-tempered Metadynamics simulations accelerate the sampling of protein-lipid conformers (Fig 2A). On the one hand, these simulations are congruent with the unbiased simulations discussed in the last paragraph, as they show a higher cholesterol binding affinity in presence of palmitoyl tails. On the other hand, we note significant differences with respect to the sampled conformational space: The Metadynamics simulations allowed us to calculate a free energy profile as a function of the coordination number. For the palmitoylated system we find a free energy minimum for the coordination around ∼70.0 (a.u.), which has not been sampled during the unbiased simulations at all (compare Fig. 1E and Fig. 2B). This reflects the sometimes major sampling and convergence difficulties for protein-lipid systems, with even 1µs long simulations being insufficient to reach the global minimum. Such problems are present in particular for non-homogeneous lipid systems.

**Figure 2:**
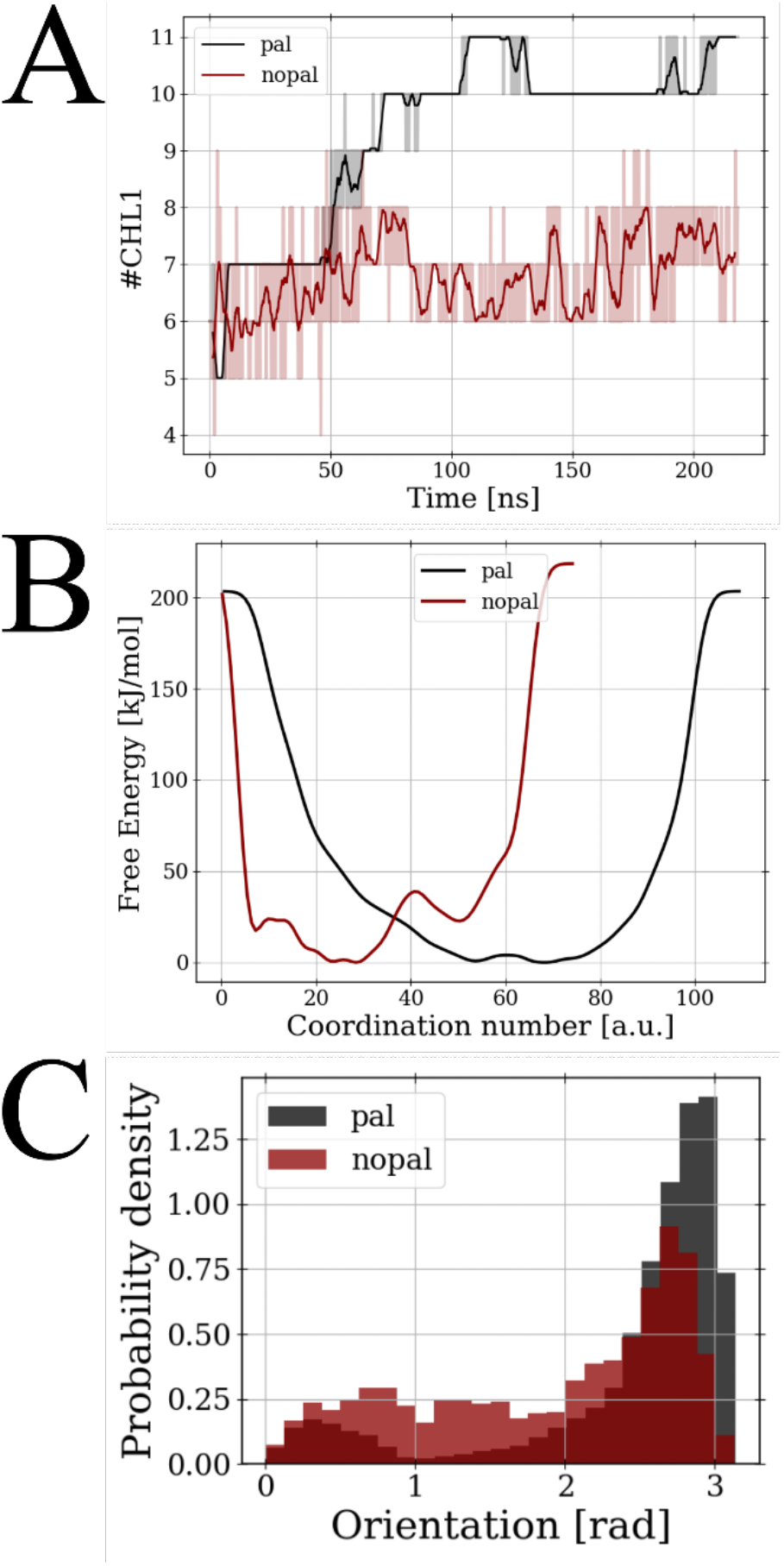
Number of cholesterol molecules in the central beta barrel during the Metadynamics simulation (A). The calculated free energy profile shows a palmitoylation-dependent shift of the global minimum (B), and different orientation distributions for the cholesterol molecules are found (C). Orientations were computed as the angle between z-axis and the vectors between hydrocarbon sidechains and the hydroxyl end. Only molecules within a cylindrical surface of 1.5 nm radius were considered.

However, we argue that our Metadynamics setup represents an effective methodology to mitigate these difficulties. The number of bound cholesterols increases rapidly during the first 100ns and fluctuates around a constant level for the second half of the simulation, which is also reflected in convergence of the free energy profiles (Fig S1,S2 of the Supporting Information). The PMFs show free energy minima at a significantly higher coordination number for the palmitoylated system, hence demonstrating a strong cholesterol-coordination effect by the palmitoyl tails. We not only monitor binding of a larger number of cholesterol molecules, but also an impact of the palmitoyl tails on the orientation of cholesterol molecules. In the non-palmitoylated system, the orientational distribution of the cholesterol is heterogeneous with a maximum around ∼ 3 rad corresponding to a vertically stacked conformation in the lipid bilayer with solvent-exposed hydroxyl groups (Fig. 2C). Population of this conformation is substantially increased, whereas inversely stacked or lateral conformations are suppressed by palmitoyl tails. Note that a more vertical stacking of cholesterols as induced by palmitoylation allows for a denser packing of cholesterol molecules, hence facilitating the accumulation of more cholesterol molecules. We hypothesize that the packing effect may arise from palmitoyl tails being very similar in length as cholesterol. From the experimental side, the role of palmitoyl tails in the CAV1-8S complex has remained unclear. However, for G-protein coupled cell receptors (GPCRs), for instance, interactions between palmitoyl and cholesterol have been shown to impact receptor signaling and other biological functions (53).

In addition, the Metadynamics simulations show intriguing transitions not sampled in the unbiased MD simulations: The CAV1-8S complex extracts individual cholesterol molecules from the distal leaflet and accommodates them in its beta-barrel (Fig 3). For the palmitoylated system we have found extraction of a single, but for the non-palmitoylated system up to three cholesterol molecules. Note that transient increases in the center of mass distances between CG atom-types of LEU174 residues of CAV1-8S and O3 cholesterol atom are attributable to orientational fluctuations of the extracted cholesterols (Fig 3B). In the palmitoylated system, the extracted cholesterol molecule mostly adopts an orientation to expose its hydroxyl group to the solvent (corresponds to 0.0 rad), whereas we note higher fluctuations in the unmodified system. The lower extraction affinity for the palmitoylated system may be explained by the more favorable stacking of cholesterols below the barrel. We stress that the extractions are stable, as reintegration of extracted cholesterols has not occurred. To the best of our knowledge, such an extraction mechanism has not yet been measured or predicted but may vitally contribute to the functioning of the CAV1-8S complex. The extraction increases the capability of the complex to accumulate cholesterol, may stabilize the central beta barrel, and slightly decreases the number of lipids in the distal leaflet. The latter effect can result in bending with positive gaussian curvature, though we expect it to be small. Kenworthy et al have hypothesized that lipids may insert into the beta-barrel (54,55). Here, we emphasize that the beta-barrel may specifically filter cholesterol, as larger lipids such as POPC are sterically and entropically unfavorable. This mechanism furthermore seems plausible, as the hydrophobic sidechains of the barrel are pointing inward, and polar or charges sidechains are solvent exposed, pointing outward (Fig. S3). Thus, the beta-barrel forms a hydrophobic pore protruding from the outer membrane layer. We also note that the barrel’s **K**I**F**SN**V** sequences (165 to 170, critical elements highlighted in bold) represent CARC domains, which are known as cholesterol binding sites in trans-membrane domains (56). From a more global perspective, we have not observed membrane-remodeling by CAV1-8S. The distal leaflet is not uniformly deformed, but most strongly bent at the position of the protein complex (Fig. 4). In addition, positive gaussian curvature occurs also exclusively at this site (FIG S4-5), and similar bending behavior is obtained for the palmitoylated and unmodified system.

**Figure 3:**
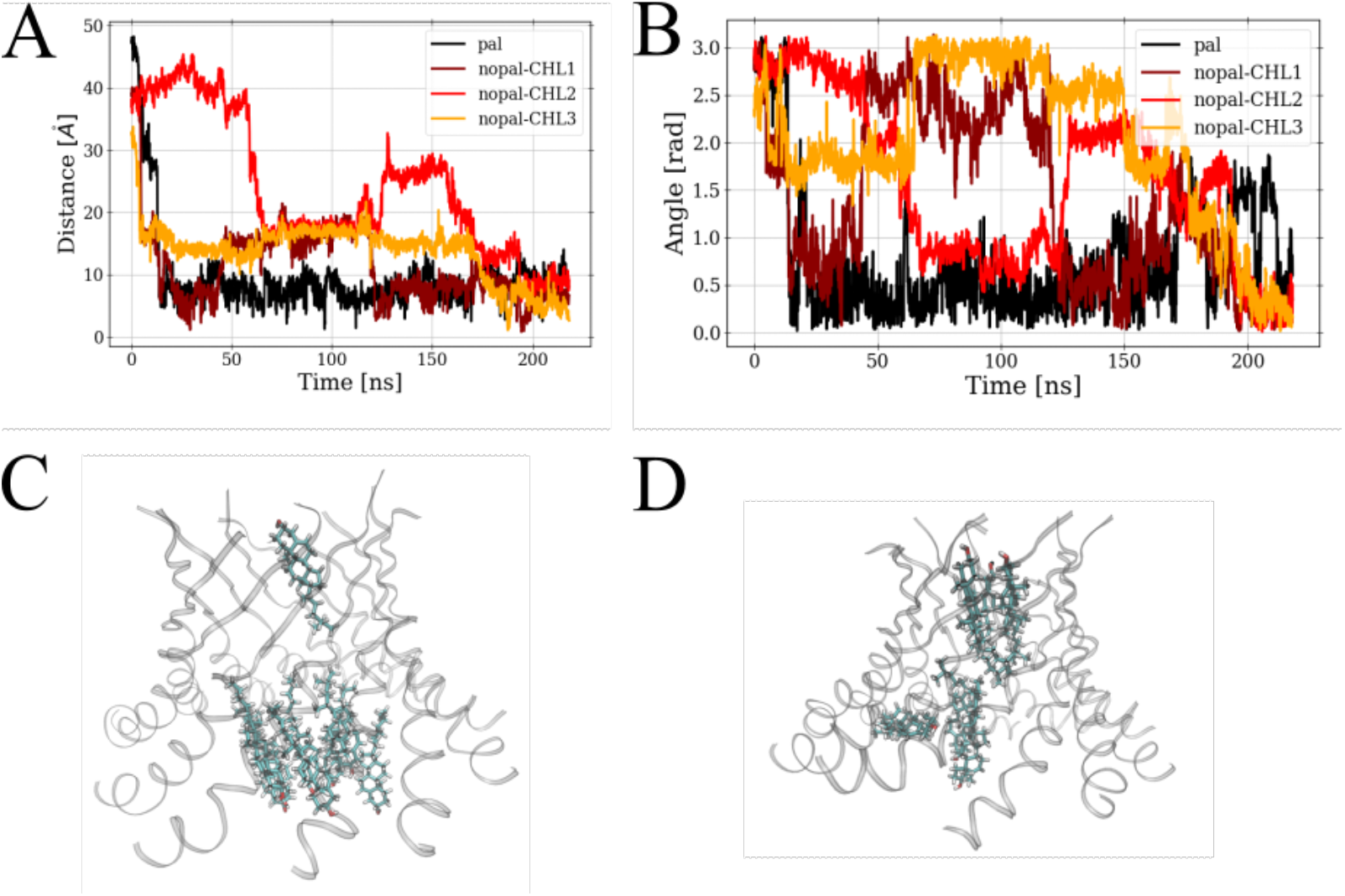
Center of mass distances between oxygen atoms of extracted cholesterol molecules and CG atom-types of LEU174 of the beta barrel (A). Orientation of extracted molecules (3.0 rad indicates hydrocarbon sidechain, and 0.0 rad indicates hydroxyl group points toward solvent- exposed C-terminal end of beta barrel) (B). Characteristic snapshot for palmitoylated and non- palmitoylated system (C,D).

**Figure 4:**
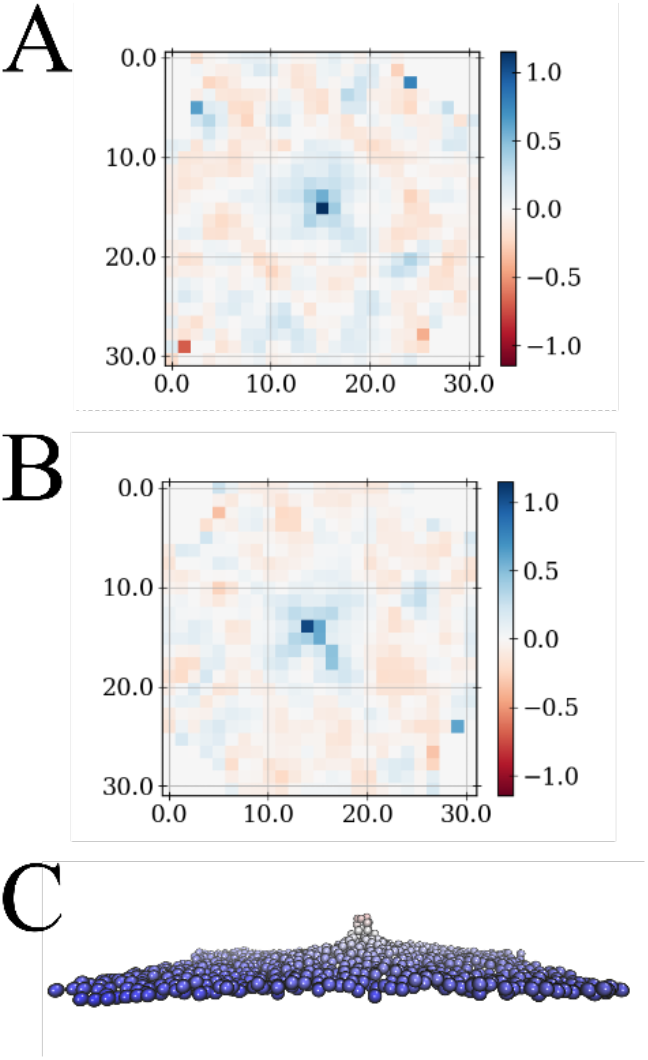
Mean-curvature computed over the distal-leaflet (25×25 pixels) for the palmitoylated and non-palmitoylated system (A,B). Characteristic snapshot of the distal-leaftlet configuration, color-coded (blue: low protrusion, red: high protrusion) in (C). The central peak is at the center of the CAV1-8S protein (omitted for clarity).

Thus, our simulations indicate a scaffolding mechanism by a single CAV1-8S, with membrane curvature being localized at the distal leaflet below the central beta barrel. Intriguingly, the mean-curvature plots indicate radial-symmetric behavior with respect to the CAV1-8S center. These fluctuations are coherent with an undulation spectrum bound by maximal undulation at the CAV1-8S beta barrel, and presence of another CAV1-8S complex may lead to interference effects. Altogether, we argue that the caveolin flat-to-curved transition likely arises as a cooperative effect between multiple CAV1-8S complexes, possibly triggered by various effects, such as: accumulation of cholesterol, interference of the undulation spectra, and electrostatic repulsion between outer rims of distinct CAV1-8S complexes (55,57). Any degree of local curvature generation is not the same as these more cooperative effects, nor does it generate the same larger scale outcome, as noted for example in the behavior of N-BAR domain proteins (58-60).

### Atomistic Simulations starting from a Martini-backmapped structure

We have noted that simulations of the CAV1-8S system with the Martini v2 CG model indicate substantially larger bending than previously discussed (61). Thus, our atomistic simulations either still suffer from a timescale problem or Martini leads to exaggerated deformations. We have tackled this enigma by backmapping the Martini structure to an atomistic structure and performing a subsequent atomistic MD simulation. Note that details on the backmapping procedure and system preparation are given in the methods section, and the simulation has been performed under unbiased conditions (i.e., no collective variable has been biased).

During this simulation, bending of the membrane reduces substantially (Fig 5), hence relaxing back to a membrane architecture similar to the previous, atomistic simulations. We emphasize that the CAV1-8S system is an oligomer, and for these systems deficiencies within Martini have been reported earlier (62). In fact, the Martini model does not maintain the secondary structure of the CAV1-8S complex, as the beta-barrel has visibly degraded. However, in our simulation we note back-relaxation only for the membrane not for the protein. As shown by Kim et al recently, exaggerated membrane deformation in Martini can occur due to insufficient barostating/neighbor-updating (63). To rule this scenario out, we have performed a Martini simulation under strict simulation-settings (see Methods, Supplement), but obtain similar results (Fig S6). Thus, we argue that the strong membrane-deformation as predicted by Martini results from inaccuracies of the entropy-enthalpy decomposition of protein-lipid interactions (29). Moreover, our backmapped-simulation also supports our previous conclusion of a membrane-scaffolding mechanism of a single CAV1-8S complex instead of global membrane remodeling.

**Figure 5:**
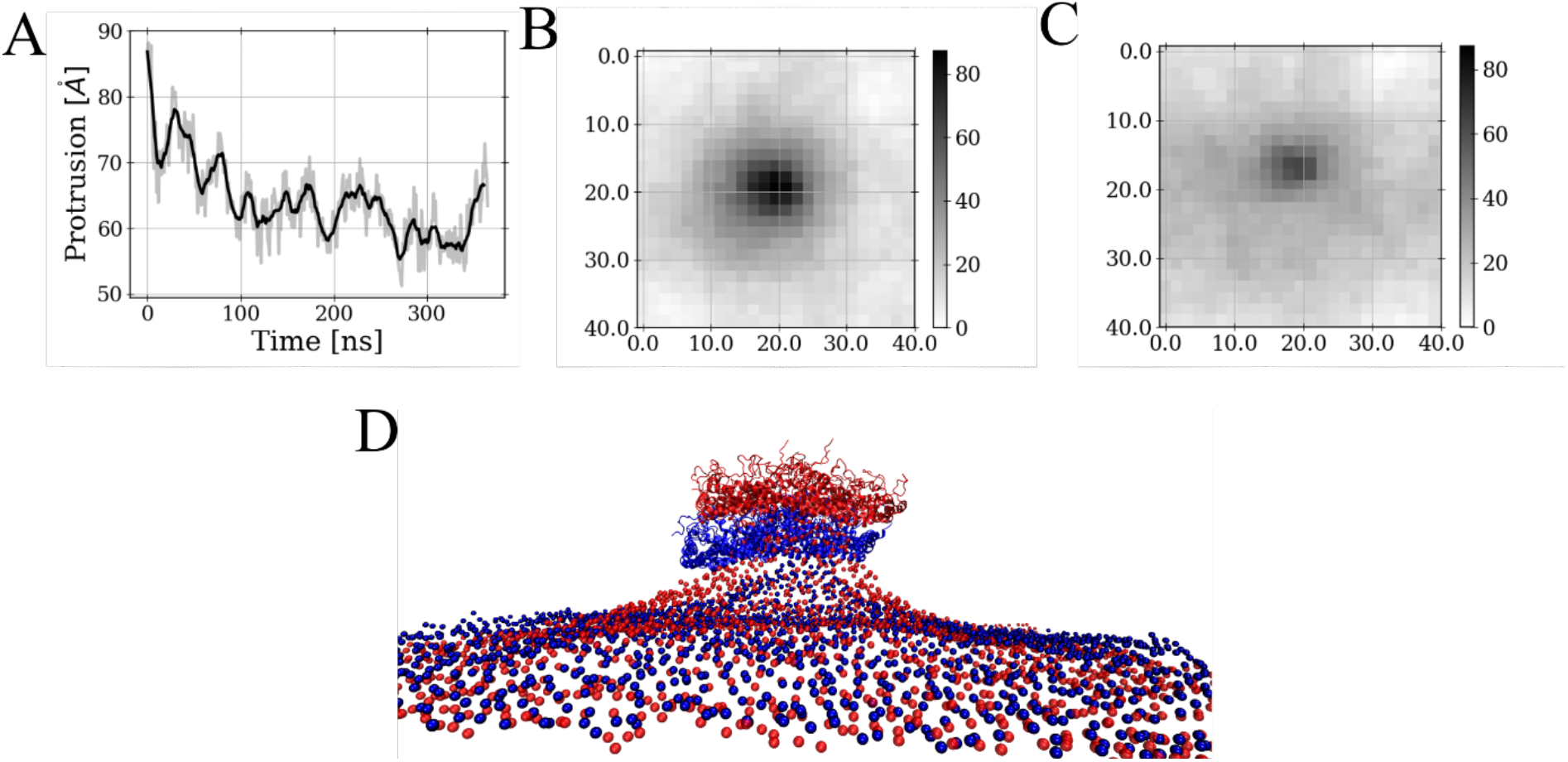
Simulation starting from Martini-backmapped structure shows substantial relaxation of the protrusion (height of protein relative to lower leaflet) (A). Height in z-direction computed over the distal-leaflet (25×25 pixels) for the starting (backmapped structure) and the final simulation snapshot, revealing major reduction of membrane undulation (B,C). Starting (red) and final (blue) snapshot of backmapped simulation. The structures were superposed on the lower leaflet, the upper leaflet is omittedforclarity. The protein has descended due to the reduction in membrane bending (D).

## Discussion

Functioning of the CAV1-8S complex has remained enigmatic for a significant period of time. Recent resolution of the complex structure has enabled us to address here some essential questions, such as cholesterol accumulation and the role of palmitoylation, with atomistic MD simulations. Such simulations can be performed on the few microsecond timescale for systems that large. We find the MD simulations to maintain the secondary structure of the complex reasonably well, but capturing the protein-lipid interactions sufficiently is still challenging due to the low diffusivity of lipids. In this paper, we also demonstrate that this obstacle can be overcome effectively with well-tempered Metadynamics simulations that employs a coordination between lipids and protein atoms as a collective variable. With this setup, we explore new and relevant conformations within ∼ 100ns of biased MD. From these simulations, we predict that the beta-barrel of the CAV1-8S system facilitates the extraction and accommodation of cholesterol molecules. This observation, to the best of our knowledge, goes beyond previous hypotheses and remains to be confirmed experimentally. Nevertheless, we argue that this mechanism is plausible due to hydrophobic interactions within the barrel. Moreover, this behavior may not only accumulate cholesterol but also impact global membrane remodeling and stabilize the structure of the beta barrel. We note that this mechanism occurred for both the palmitoylated and unmodified system. This speaks in favor of cholesterol accumulation occurring also in the absence of palmitoylation and may serve as an explanation why caveolae maturation has also been reported when palmitoylation was eliminated [4]. Quantitatively, however, we find a cholesterol stabilizing role of palmitoylations. The number of cholesterols in proximity to the barrel is larger in the presence of palmitoylations, which organize a vertical stacking pattern of the cholesterol molecules. As a consequence, palmitoylation should at least increase the affinity for caveolae maturation.

We furthermore find qualitatively the same membrane deformation for both systems. The MD simulations do not exhibit global membrane bending, but instead localized curvature at the position of the beta barrel. This scaffolding mechanism is further supported by atomistic MD simulations starting from a significantly deformed structure sampled with the Martini 2 CG model, as bending relaxes substantially in the backmapped Martini to all-atom simulation. The Martini 2 CG and atomistic MD simulations therefore lead to different interpretations. While the former indicates membrane remodeling already occurring as a single-complex phenomenon, atomistic MD simulations indicate that the flat-to-curved transition is likely to be caused cooperatively between multiple CAV1-8S complexes. We find the atomistic MD results to also be more in line with the dodecahedral models elaborated from experiments. Further testing of this notion requires the development of a multiscale CG model probably of coarser resolution than Martini. This will be addressed in the near future, as our atomistic MD simulations provide the required “bottom-up” reference data for such coarse-graining.

## Conclusions

We conclude that palmitoylation may not be required for cholesterol accumulation by the CAV1-8S complex, but at least enhances this effect. It generates an ordered, stacked arrangement of cholesterol molecules, but the global shape of the membrane remains similar to the unmodified system. Here, our simulations also indicate that a single CAV1-8S complex does not remodel the membrane structure, so multiple complexes may be required to cooperatively induce a flat-to-curved transition. From the methodological point of view, we furthermore conclude that well-tempered Metadynamics simulations biasing protein-lipid coordination represents an effective approach that can also be used for other protein-membrane systems. Using this setup, the trajectories readily visit the free energy minimum that has not been sampled in an unbiased 1 µs atomistic MD simulation and hence the Metadynamics facilitated relevant new insight into the conformational behavior of CAV1-8S. Besides, we predict that the beta barrel can serve as a reservoir to store cholesterol molecules.

In addition, we also demonstrate that backmapping from a CG model to an all-atom one can be used as a powerful tool to scrutinize coarse-grained model predictions even for larger protein-membrane systems. For the CAV1-8S complex, we find discrepancies between atomistic MD and Martini 2 CG simulations, especially with respect to the deformation of the membrane by the CAV1-8S complex.

## Acknowledgements

This work was supported by an award from the National Institute of General Medical Science (NIGMS) of the National Institutes of Health (NIH), grant R01GM063796. The content is solely the responsibility of the authors and does not necessarily represent the official views of the National Institutes of Health. This project also received funding from the European Union’s Framework Programme for Research and Innovation Horizon Europe (2021-2027) under the Marie Skłodowska-Curie Grant Agreement No. 101109916 to K.L.. Computational resources were provided by the University of Chicago Research Computing Center, the Extreme Science and Engineering Discovery Environment (XSEDE), and the NIH-funded Beagle-3 computer (NIH award 1S10OD028655).

## Bibliography

1. Root, K. T., J. A. Julien, and K. J. Glover (2019). Secondary structure of caveolins: A mini review. Biochemical Society Transactions.

2. Ohi, M. D., and A. K. Kenworthy (2022). Emerging Insights into the Molecular Architecture of Caveolin-1. Journal of Membrane Biology.

3. Porta, J. C., B. Han, A. Gulsevin, J. M. Chung, Y. Peskova, S. Connolly, H. S. McHaourab, J. Meiler, E. Karakas, A. K. Kenworthy, and M. D. Ohi. 2022. Molecular architecture of the human caveolin-1 complex. Science Advances. 8(19), doi: 10.1126/sciadv.abn7232.

4. Dietzen, D. J., W. R. Hastings, and D. M. Lublin. 1995. Caveolin is palmitoylated on multiple cysteine residues. Palmitoylation is not necessary for localization of caveolin to caveolae. Journal of Biological Chemistry. 270(12), doi: 10.1074/jbc.270.12.6838.

5. Parat, M. O., and P. L. Fox. 2001. Palmitoylation of Caveolin-1 in Endothelial Cells is Post-translational but Irreversible. Journal of Biological Chemistry. 276(19), doi: 10.1074/jbc.M006722200.

6. Hojo, H., T. Takei, Y. Asahina, N. Okumura, T. Takao, M. So, I. Suetake, T. Sato, A. Kawamoto, and Y. Hirabayashi. 2021. Total Synthesis and Structural Characterization of Caveolin-1. Angewandte Chemie - International Edition. 60(25), doi: 10.1002/anie.202100826.

7. Nørholm, M. H. H., Y. V. Shulga, S. Aoki, R. M. Epand, and G. Von Heijne. 2011. Flanking residues help determine whether a hydrophobic segment adopts a monotopic or bitopic topology in the endoplasmic reticulum membrane. Journal of Biological Chemistry. 286(28), doi: 10.1074/jbc.M111.244616.

8. Rothberg, K. G., J. E. Heuser, W. C. Donzell, Y. S. Ying, J. R. Glenney, and R. G. W. Anderson. 1992. Caveolin, a protein component of caveolae membrane coats. Cell. 68(4), doi: 10.1016/0092-8674(92)90143-Z.

9. Ludwig, A., B. J. Nichols, and S. Sandin. 2016. Architecture of the caveolar coat complex. Journal of Cell Science. 129(16), doi: 10.1242/jcs.191262.

10. Stoeber, M., P. Schellenberger, C. A. Siebert, C. Leyrat, K. Grünewald, and A. Helenius. 2016. Model for the architecture of caveolae based on a flexible, net-like assembly of Cavin1 and Caveolin discs. Proceedings of the National Academy of Sciences of the United States of America. 113(50), doi: 10.1073/pnas.1616838113.

11. Lamaze, C., N. Tardif, M. Dewulf, S. Vassilopoulos, and C. M. Blouin (2017). The caveolae dress code: structure and signaling. Current Opinion in Cell Biology.

12. Khater, I. M., Q. Liu, K. C. Chou, G. Hamarneh, and I. R. Nabi. 2019. Super-resolution modularity analysis shows polyhedral caveolin-1 oligomers combine to form scaffolds and caveolae. Scientific Reports. 9(1), doi: 10.1038/s41598-019-46174-z.

13. Parton, R. G., and M. A. Del Pozo. 2013. Caveolae as plasma membrane sensors, protectors and organizers. Nature Reviews Molecular Cell Biology. 14(2), doi: 10.1038/nrm3512.

14. Lolo, F. N., N. Walani, E. Seemann, D. Zalvidea, D. M. Pavón, G. Cojoc, M. Zamai, C. Viaris de Lesegno, F. Martínez de Benito, M. Sánchez-Álvarez, J. J. Uriarte, A. Echarri, D. Jiménez-Carretero, J. C. Escolano, S. A. Sánchez, V. R. Caiolfa, D. Navajas, X. Trepat, J. Guck, C. Lamaze, P. Roca-Cusachs, M. M. Kessels, B. Qualmann, M. Arroyo, and M. A. del Pozo. 2023. Caveolin-1 dolines form a distinct and rapid caveolae-independent mechanoadaptation system. Nature Cell Biology. 25(1), doi: 10.1038/s41556-022-01034-3.

15. Frank, P. G., S. Pavlides, M. W. C. Cheung, K. Daumer, and M. P. Lisanti. 2008. Role of caveolin-1 in the regulation of lipoprotein metabolism. American Journal of Physiology - Cell Physiology. 295(1), doi: 10.1152/ajpcell.00185.2008.

16. Fridolfsson, H. N., D. M. Roth, P. A. Insel, and H. H. Patel (2014). Regulation of intracellular signaling and function by caveolin. FASEB Journal.

17. Han, B., C. A. Copeland, Y. Kawano, E. B. Rosenzweig, E. D. Austin, L. Shahmirzadi, S. Tang, K. Raghunathan, W. K. Chung, and A. K. Kenworthy. 2016. Characterization of a caveolin-1 mutation associated with both pulmonary arterial hypertension and congenital generalized lipodystrophy. Traffic. 17(12), doi: 10.1111/tra.12452.

18. Tian, J., M. S. Popal, R. C. Huang, M. Zhang, X. Zhao, M. Zhang, and X. Song (2020). Caveolin as a novel potential therapeutic target in cardiac and vascular diseases: A mini review. Aging and Disease.

19. Williams, T. M., and M. P. Lisanti (2005). Caveolin-1 in oncogenic transformation, cancer, and metastasis. American Journal of Physiology - Cell Physiology.

20. Gupta, R., C. Toufaily, and B. Annabi (2014). Caveolin and cavin family members: Dual roles in cancer. Biochimie.

21. Frank, P. G., Y. L. Marcel, M. A. Connelly, D. M. Lublin, V. Franklin, D. L. Williams, and M. P. Lisanti. 2002. Stabilization of caveolin-1 by cellular cholesterol and scavenger receptor class B type I. Biochemistry. 41(39), doi: 10.1021/bi0257078.

22. Koh, S., W. Lee, S. M. Park, and S. H. Kim. 2021. Caveolin-1 deficiency impairs synaptic transmission in hippocampal neurons. Molecular Brain. 14(1), doi: 10.1186/s13041-021-00764-z.

23. Tillu, V. A., J. Rae, Y. Gao, N. Ariotti, M. Floetenmeyer, O. Kovtun, K. A. McMahon, N. Chaudhary, R. G. Parton, and B. M. Collins. 2021. Cavin1 intrinsically disordered domains are essential for fuzzy electrostatic interactions and caveola formation. Nature Communications. 12(1), doi: 10.1038/s41467-021-21035-4.

24. Kozlov, M. M., and J. W. Taraska (2023). Generation of nanoscopic membrane curvature for membrane trafficking. Nature Reviews Molecular Cell Biology.

25. Krishna, A., and D. Sengupta. 2019. Interplay between Membrane Curvature and Cholesterol: Role of Palmitoylated Caveolin-1. Biophysical Journal. 116(1), doi: 10.1016/j.bpj.2018.11.3127.

26. Prakash, S., H. Malshikare, and D. Sengupta (2022). Molecular Mechanisms Underlying Caveolin-1 Mediated Membrane Curvature. Journal of Membrane Biology.

27. Barducci, A., G. Bussi, and M. Parrinello. 2008. Well-tempered metadynamics: A smoothly converging and tunable free-energy method. Physical Review Letters. 100(2), doi: 10.1103/PhysRevLett.100.020603.

28. Dama, J. F., M. Parrinello, and G. A. Voth. 2014. Well-Tempered Metadynamics Converges Asymptotically. Physical Review Letters. 112(24):240602, doi: 10.1103/PhysRevLett.112.240602, https://link.aps.org/doi/10.1103/PhysRevLett.112.240602.

29. Jarin, Z., J. Newhouse, and G. A. Voth. 2021. Coarse-Grained Force Fields from the Perspective of Statistical Mechanics: Better Understanding of the Origins of a MARTINI Hangover. Journal of Chemical Theory and Computation. 17(2), doi: 10.1021/acs.jctc.0c00638.

30. Lomize, M. A., I. D. Pogozheva, H. Joo, H. I. Mosberg, and A. L. Lomize. 2012. OPM database and PPM web server: Resources for positioning of proteins in membranes. Nucleic Acids Research. 40(D1), doi: 10.1093/nar/gkr703.

31. Jo, S., T. Kim, V. G. Iyer, and W. Im. 2008. CHARMM-GUI: A web-based graphical user interface for CHARMM. Journal of Computational Chemistry. 29(11), doi: 10.1002/jcc.20945.

32. Lee, J., X. Cheng, J. M. Swails, M. S. Yeom, P. K. Eastman, J. A. Lemkul, S. Wei, J. Buckner, J. C. Jeong, Y. Qi, S. Jo, V. S. Pande, D. A. Case, C. L. Brooks, A. D. MacKerell, J. B. Klauda, and W. Im. 2016. CHARMM-GUI Input Generator for NAMD, GROMACS, AMBER, OpenMM, and CHARMM/OpenMM Simulations Using the CHARMM36 Additive Force Field. Journal of Chemical Theory and Computation. 12(1), doi: 10.1021/acs.jctc.5b00935.

33. Huang, J., S. Rauscher, G. Nawrocki, T. Ran, M. Feig, B. L. De Groot, H. Grubmüller, and A. D. MacKerell. 2016. CHARMM36m: An improved force field for folded and intrinsically disordered proteins. Nature Methods. 14(1), doi: 10.1038/nmeth.4067.

34. Jorgensen, W. L., J. Chandrasekhar, J. D. Madura, R. W. Impey, and M. L. Klein. 1983. Comparison of simple potential functions for simulating liquid water. The Journal of Chemical Physics. 79(2), doi: 10.1063/1.445869.

35. Van Der Spoel, D., E. Lindahl, B. Hess, G. Groenhof, A. E. Mark, and H. J. C. Berendsen (2005). GROMACS: Fast, flexible, and free. Journal of Computational Chemistry.

36. Berendsen, H. J. C., J. P. M. Postma, W. F. Van Gunsteren, A. Dinola, and J. R. Haak. 1984. Molecular dynamics with coupling to an external bath. The Journal of Chemical Physics. 81(8), doi: 10.1063/1.448118.

37. Parrinello, M., and A. Rahman. 1981. Polymorphic transitions in single crystals: A new molecular dynamics method. Journal of Applied Physics. 52(12), doi: 10.1063/1.328693.

38. Nosé, S. 1984. A unified formulation of the constant temperature molecular dynamics methods. The Journal of Chemical Physics. 81(1), doi: 10.1063/1.447334.

39. Hoover, W. G. 1985. Canonical dynamics: Equilibrium phase-space distributions. Physical Review A. 31(3), doi: 10.1103/PhysRevA.31.1695.

40. Humphrey, W., A. Dalke, and K. Schulten. 1996. VMD: Visual molecular dynamics. Journal of Molecular Graphics. 14(1), doi: 10.1016/0263-7855(96)00018-5.

41. Michaud-Agrawal, N., E. J. Denning, T. B. Woolf, and O. Beckstein. 2011. MDAnalysis: A toolkit for the analysis of molecular dynamics simulations. Journal of Computational Chemistry. 32(10), doi: 10.1002/jcc.21787.

42. Gowers, R., M. Linke, J. Barnoud, T. Reddy, M. Melo, S. Seyler, J. Domański, D. Dotson, S. Buchoux, I. Kenney, and O. Beckstein (2016). MDAnalysis: A Python Package for the Rapid Analysis of Molecular Dynamics Simulations. Proceedings of the 15th Python in Science Conference.

43. Laio, A., and M. Parrinello. 2002. Escaping free-energy minima. Proceedings of the National Academy of Sciences of the United States of America. 99(20), doi: 10.1073/pnas.202427399.

44. Tribello, G. A., M. Bonomi, D. Branduardi, C. Camilloni, and G. Bussi. 2014. PLUMED 2: New feathers for an old bird. Computer Physics Communications. 185(2), doi: 10.1016/j.cpc.2013.09.018.

45. Qi, Y., H. I. Ingólfsson, X. Cheng, J. Lee, S. J. Marrink, and W. Im. 2015. CHARMM-GUI Martini Maker for Coarse-Grained Simulations with the Martini Force Field. Journal of Chemical Theory and Computation. 11(9), doi: 10.1021/acs.jctc.5b00513.

46. Hsu, P. C., B. M. H. Bruininks, D. Jefferies, P. Cesar Telles de Souza, J. Lee, D. S. Patel, S. J. Marrink, Y. Qi, S. Khalid, and W. Im. 2017. Charmm-gui martini maker for modeling and simulation of complex bacterial membranes with lipopolysaccharides. Journal of Computational Chemistry. 38(27), doi: 10.1002/jcc.24895.

47. De Jong, D. H., G. Singh, W. F. D. Bennett, C. Arnarez, T. A. Wassenaar, L. V. Schäfer, X. Periole, D. P. Tieleman, and S. J. Marrink. 2013. Improved parameters for the martini coarse-grained protein force field. Journal of Chemical Theory and Computation. 9(1), doi: 10.1021/ct300646g.

48. Souza, P. C. T., R. Alessandri, J. Barnoud, S. Thallmair, I. Faustino, F. Grünewald, I. Patmanidis, H. Abdizadeh, B. M. H. Bruininks, T. A. Wassenaar, P. C. Kroon, J. Melcr, V. Nieto, V. Corradi, H. M. Khan, J. Domański, M. Javanainen, H. Martinez-Seara, N. Reuter, R. B. Best, I. Vattulainen, L. Monticelli, X. Periole, D. P. Tieleman, A. H. de Vries, and S. J. Marrink. 2021. Martini 3: a general purpose force field for coarse-grained molecular dynamics. Nature Methods. 18(4):382–388, doi: 10.1038/s41592-021-01098-3, 10.1038/s41592-021-01098-3.

49. Borges-Araújo, L., A. C. Borges-Araújo, T. N. Ozturk, D. P. Ramirez-Echemendia, B. Fábián, T. S. Carpenter, S. Thallmair, J. Barnoud, H. I. Ingólfsson, G. Hummer, D. P. Tieleman, S. J. Marrink, P. C. T. Souza, and M. N. Melo. 2023. Martini 3 Coarse-Grained Force Field for Cholesterol. Journal of Chemical Theory and Computation. 19(20):7387–7404, doi: 10.1021/acs.jctc.3c00547, 10.1021/acs.jctc.3c00547.

50. Bussi, G., D. Donadio, and M. Parrinello. 2007. Canonical sampling through velocity rescaling. Journal of Chemical Physics. 126(1), doi: 10.1063/1.2408420.

51. Wassenaar, T. A., K. Pluhackova, R. A. Böckmann, S. J. Marrink, and D. P. Tieleman. 2014. Going backward: A flexible geometric approach to reverse transformation from coarse grained to atomistic models. Journal of Chemical Theory and Computation. 10(2), doi: 10.1021/ct400617g.

52. Kim, S. 2023. Backmapping with Mapping and Isomeric Information. The Journal of Physical Chemistry B. 127(49):10488–10497, doi: 10.1021/acs.jpcb.3c05593, 10.1021/acs.jpcb.3c05593.

53. Chini, B., and M. Parenti (2009). G-protein-coupled receptors, cholesterol and palmitoylation: Facts about fats. Journal of Molecular Endocrinology.

54. Kenworthy, A. K. 2023. The building blocks of caveolae revealed: caveolins finally take center stage. Biochemical Society Transactions. 51(2):855–869.

55. Kenworthy, A. K., B. Han, N. Ariotti, and R. G. Parton. 2023. The role of membrane lipids in the formation and function of caveolae. Cold Spring Harbor Perspectives in Biology. 15(9):a041413.

56. Fantini, J., and F. J. Barrantes (2013). How cholesterol interacts with membrane proteins: An exploration of cholesterol-binding sites including CRAC, CARC, and tilted domains. Frontiers in Physiology.

57. Simunovic, M., A. Srivastava, and G. A. Voth. 2013. Linear aggregation of proteins on the membrane as a prelude to membrane remodeling. Proceedings of the National Academy of Sciences. 110(51):20396–20401, doi: 10.1073/pnas.1309819110, 10.1073/pnas.1309819110.

58. Ayton, G. S., P. D. Blood, and G. A. Voth. 2007. Membrane remodeling from N-BAR domain interactions: insights from multi-scale simulation. Biophysical journal. 92(10):3595–3602.

59. Blood, P. D., R. D. Swenson, and G. A. Voth. 2008. Factors influencing local membrane curvature induction by N-BAR domains as revealed by molecular dynamics simulations. Biophysical journal. 95(4):1866–1876.

60. Simunovic, M., C. Mim, T. C. Marlovits, G. Resch, V. M. Unger, and G. A. Voth. 2013. Protein-mediated transformation of lipid vesicles into tubular networks. Biophysical journal. 105(3):711–719.

61. Milka Doktorova, A. K. I. L. private communication.

62. Javanainen, M., H. Martinez-Seara, and I. Vattulainen. 2017. Excessive aggregation of membrane proteins in the Martini model. PLoS ONE. 12(11), doi: 10.1371/journal.pone.0187936.

63. Kim, H., B. Fábián, and G. Hummer. 2023. Neighbor List Artifacts in Molecular Dynamics Simulations.

