## Supporting Information for "Lipid Organization by the Caveolin-1 Complex"

**Corresponding Author**:

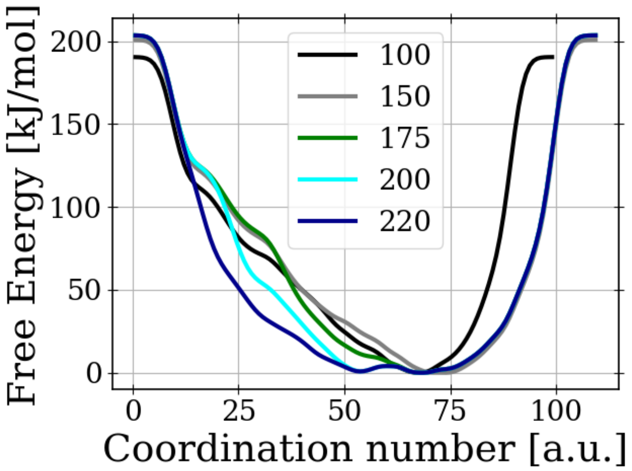


SI FIG S1: Computed Free energy profiles for fractions of the simulation (in ns), for the palmitoylated system.

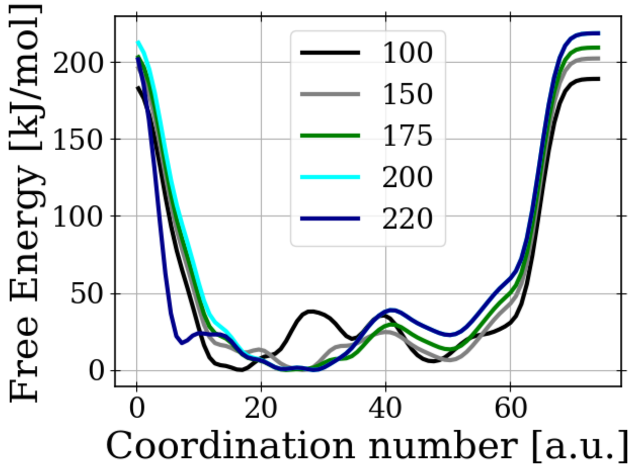


SI FIG S2: Computed Free energy profiles for fractions of the simulation (in ns), for the non-palmitoylated system.

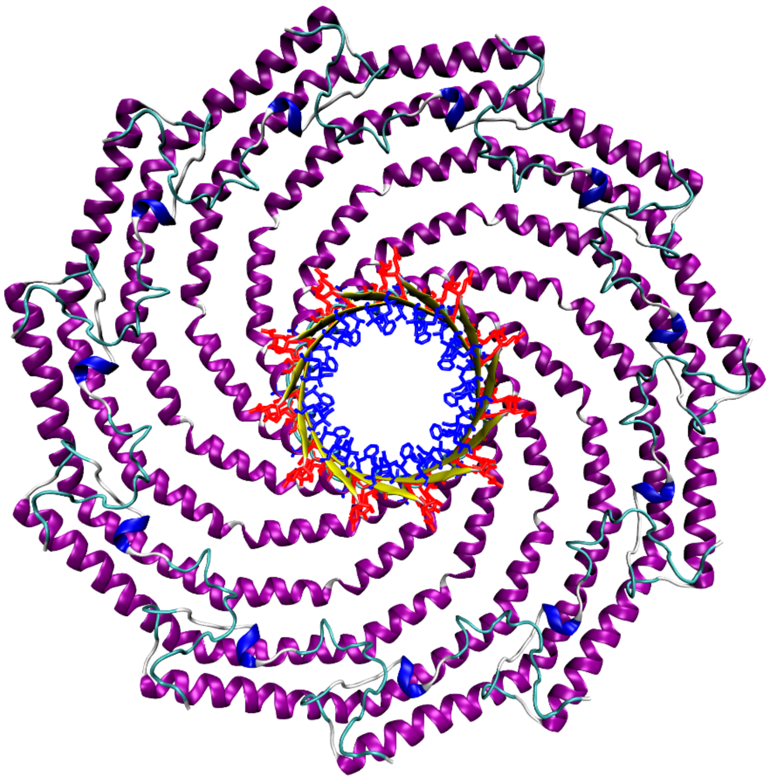

SI FIG S3: Structure of the CAV1-8S complex. Hydrophobic residues of the beta barrel are highlighted as blue, polar or charged residues as red.

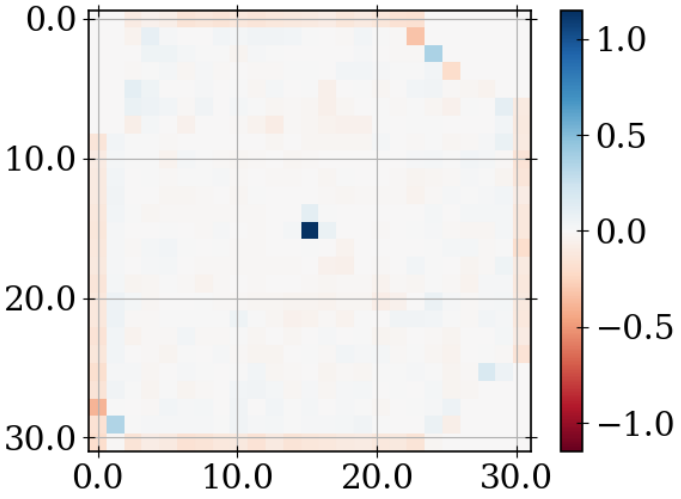


FIG S4: Average Gaussian curvature computed for the lower leaflet of the palmitoylated system (from Metadynamics simulation).

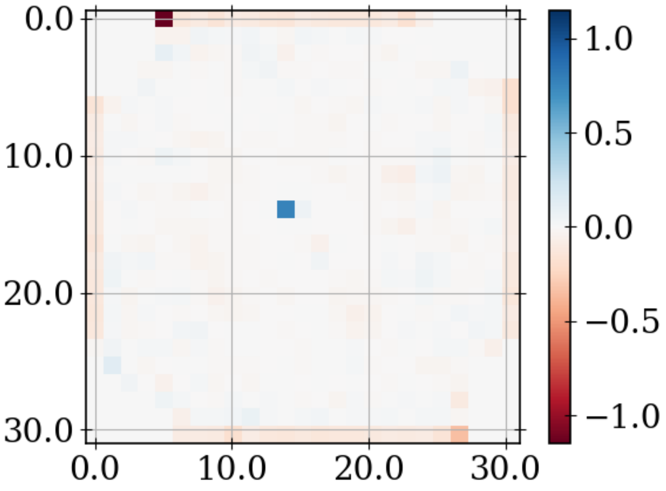


SI FIG S5: Average Gaussian curvature computed for the lower leaflet of the non-palmitoylated system (from Metadynamics simulation).

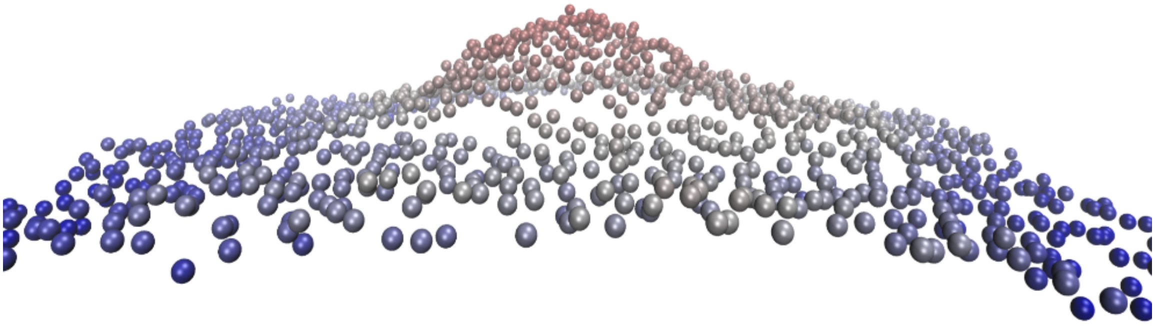

SI FIG 6: Lower leaflet sampled from 100ns Martini simulation. PO4 beads are shown color coded, blue low z-value, red high z-value.
